# Novel ionophores active against La Crosse virus identified through rapid antiviral screening

**DOI:** 10.1101/2020.01.21.914929

**Authors:** Zachary J. Sandler, Michelle N. Vu, Vineet D. Menachery, Bryan C. Mounce

## Abstract

Bunyaviruses are significant human pathogens, causing diseases ranging from hemorrhagic fevers to encephalitis. Among these viruses, La Crosse virus (LACV), a member of the California serogroup, circulates in the eastern and midwestern United States. While LACV infection is often asymptomatic, dozens of cases of encephalitis are reported yearly. Unfortunately, no antivirals have been approved to treat LACV infection. Here, we developed a method to rapidly test potential antivirals against LACV infection. From this screen, we identified several potential antiviral molecules, including known antivirals. Additionally, we identified many novel antivirals that exhibited antiviral activity without affecting cellular viability. Valinomycin, a potassium ionophore, was among our top targets. We found that valinomycin exhibited potent anti-LACV activity in multiple cell types in a dose-dependent manner. Valinomycin did not affect particle stability or infectivity, suggesting that it may preclude virus replication by altering cellular potassium ions, a known determinant of LACV entry. We extended these results to other ionophores and found that the antiviral activity of valinomycin extended to other viral families including bunyaviruses (Rift Valley fever virus, Keystone virus), enteroviruses (Coxsackievirus, rhinovirus), flavirivuses (Zika), and coronaviruses (229E and MERS-CoV). In all viral infections, we observed significant reductions in virus titer in valinomycin-treated cells. In sum, we demonstrate the importance of potassium ions to virus infection, suggesting a potential therapeutic target to disrupt virus replication.

**Importance:** No antivirals are approved for the treatment of bunyavirus infection. The ability to rapidly screen compounds and identify novel antivirals is one means to accelerate drug discovery for viruses with no approved treatments. We used this approach to screen hundreds of compounds against La Crosse virus, an emerging bunyavirus that causes significant disease, including encephalitis. We identified several known and previously unidentified antivirals. We focused on a potassium ionophore, valinomycin, due to its promising *in vitro* antiviral activity. We demonstrate that valinomycin, as well as a selection of other ionophores, exhibits activity against La Crosse virus as well as several other distantly related bunyaviruses. We finally observe that valinomycin has activity against a wide array of human viral pathogens, suggesting that disrupting potassium ion homeostasis with valinomycin may be a potent host pathway to target to quell virus infection.

## Introduction

Bunyaviruses are the largest family of viruses, composed of hundreds of members. These segmented, negative-sense RNA viruses are transmitted primarily by an arthropod vector, and several family members pose significant threats to human health. Rift Valley fever virus (RVFV) frequently infects humans and ruminants, resulting in severe morbidity and mortality. While RVFV is primarily transmitted in Africa and the Middle East, the threat of global spread looms, and recent examples of Zika^1^ and chikungunya^2^ viruses illustrate this tangible hazard. In addition to RVFV, several other bunyaviruses infect humans and are associated with severe pathologies as well. La Crosse virus (LACV), distantly related to RVFV, is a bunyavirus present primarily in the midwestern and eastern United States^3^. LACV is transmitted by *Aedes triseriatus*, though *Aedes albopictus*^4–6^ efficiently carries the virus as well. While relatively unknown and frequently undiagnosed, LACV infects and causes neuroinvasive disease in dozens of people every year. In fact, from 2009 to 2018, 679 such cases were reported according to the Centers for Disease Control^7^. Further, LACV cases recently have emerged in the southeastern United States, suggesting spread of the virus^8^. The ability of LACV to infect *Aedes albopictus* mosquitos will likely lead to its continuing spread. Despite this spread and the severe disease associated with LACV infection, no antivirals or vaccines are available to prevent or treat infection. Thus, LACV presents a significant threat to human health.

Despite their prevalence, no antivirals are available to treat bunyavirus infection and palliative care is given to patients presenting with LACV encephalitis. While several vaccine candidates have been developed to target RVFV^9–11^, no similar effort has been invested in anti-LACV therapeutics. With the increasing availability of drug panels to screen molecules for antivirals, rapid investigation and deployment of antivirals for emerging viruses is possible. Several prior screens for antivirals have successfully identified lead molecules and highlighted cellular pathways critical to virus infection. For example, recent screens with chikungunya virus highlighted berberine, abamectin, and ivermectin as promising antivirals^12^. Additionally, Zika virus screens have uncovered known and novel antivirals^13–15^, including mycophenolic acid and daptomycin, among others. More closely related to LACV, a RVFV screen highlighted azauridine and mitoxatrone^16^. The ability to rapidly screen molecules highlights an opportunity to identify both unique virus-specific and broad-spectrum antivirals, as these reports have highlighted.

Antivirals may target viral particles, viral processes and critical host-targeted antivirals^17^. Host-directed antivirals have gained appreciation recently, as interfering with host processes crucial to viral processes has several benefits, including potential for broad-spectrum activity and a heightened requirement for antiviral resistance beyond minor mutations in the virus. Additionally, numerous approved and available drugs are already available that can be repurposed as antivirals^18,19^. While significant work remains to be done to identify and verify such antivirals, rapidly screening compounds can provide insight.

Using the NIH’s Developmental Therapeutics Program (DTP), we obtained and screened >500 compounds for activity against LACV. We identified several known antivirals, including deoxyuridine and quinonone. Importantly, we also identified a variety of novel classes of antivirals, including metal ion chelators. Valinomycin, a top hit in our screen, functions by transporting potassium ions against the electrochemical gradient. We investigated the antiviral activity of valinomycin, observing that valinomycin exhibits antiviral activity in several cellular systems, in a dose-dependent manner and independent of treatment time. We also found that valinomycin does not directly inactivate viral particles, highlighting a cellular role for potassium ions in virus infection. We expanded our results to additional ionophores, observing that some but not all effectively blocked LACV replication. Finally, we determined that valinomycin is broadly antiviral, as it reduced replication of several viruses from diverse families, including flaviviruses and enteroviruses. Together, these data highlight the utility in rapid screening of antiviral molecules as well as a crucial role for potassium ions in LACV infection.

## Results

### Development of rapid screening of NIH DTP compounds active against LACV

We developed a simple, rapid assay to measure antiviral activity in Huh7 cells (Figure 1A). We plated Huh7 cells to confluency in 96-well plates, to which we added 2 µM drug from the NIH NCI Development Therapeutics Program (DTP). Two hours late, cells were infected at multiplicity of infection (MOI) of 0.1 plaque-forming units (pfu) per cell. At 48 h post infection (hpi), cells were fixed with formalin and stained with crystal violet stain. Because viable cells robustly stain with crystal violet, while dead cells do not, we could discriminate between live and dead cells. Importantly, any cytotoxic molecule would not stain with crystal violet; thus, stained cells indicate antiviral molecules that are not cytotoxic at 2 µM. Crystal violet stain was subsequently resuspended in 10% acetic acid and absorbance read at 595 nm. To control for inter-plate variability, each plate contained an untreated and infected (low survival), untreated and uninfected (high survival), and ribavirin-treated (400 µM) control (high survival). Absorbances of drug-treated and infected wells (Figure 1B) were compared to untreated and uninfected controls by dividing their absorbance values (Figure 1C). This ratio highlighted several candidate antivirals, including lagistase, lapachol, superacyl, and valinomycin. Interestingly, we identified several known antivirals in our screen, including deoxyuridine and nelarabine (summarized in Table 1). Thus, our assay identified molecules with novel activity against LACV, including some recognized antivirals.

**Table 1.**

| DTP NSC Number | Primary Screen Hit Value | Name | Description | Prior Antiviral Activity |
| --- | --- | --- | --- | --- |
| 407286 | 16.97 | Lagistase | Naturally occurring plant phenol; antioxidant and metabolism-modulating properties | HIV<sup>32</sup>, HRV<sup>33</sup>, HBV<sup>34</sup> |
| 11905 | 5.76 | Lapachol | Naphthoquinone from the lapacho tree; anti-inflammatory and anti-proliferation agent | EBV<sup>35</sup>, EV<sup>36</sup> |
| 41833 | 3.97 | Superacyl | Base analog |  |
| 122023 | 3.19 | Valinomycin | Potassium ionophore | HRV<sup>37</sup>, SARS-CoV<sup>26</sup>, RSV<sup>38</sup>, PV<sup>39</sup> |
| 51787 | 3.19 | 5-Isoquinolinol | Metal chelating agent |  |
| 24819 | 2.93 | β-peltatin | Glycoside from *Podophyllum* roots | MV, HSV<sup>40</sup> |
| 401005 | 2.84 | Pleurotine | Antifungal agent; thioredoxin inhibitor |  |
| 44175 | 2.59 | Polyporin | Benzoquinone isolated from *Hapalopilus nidulans*; dihydroorotate dehydrogenase inhibitor |  |
| 712807 | 2.48 | Capecitabine | Antimetabolite antineoplastic agent |  |
| 15307 | 2.45 | Quininone | Quinine metabolite |  |
| 36398 | 2.20 | Taxifolin | Polyphenol from milk thistle seeds | HAV<sup>41</sup>, HIV<sup>42</sup> |
| 38010 | 2.15 | Chaulmoogric acid ethyl ester | Ethyl ester of long chain fatty acid from chaulmoogra seeds |  |
| 747972 | 2.10 | Lenalidomide | Anti-angiogenic factor, used to treat multiple myeloma |  |
| 718781 | 2.04 | Tarceva | Epidermal growth factor receptor inhibitor antineoplastic agent, used to treat pancreatic and non-small cell lung cancer |  |
| 301683 | 2.02 | Daphnetin | Antioxidant agent; coumarin; used in traditional Chinese medicine to treat cardiovascular disease |  |
| 755985 | 1.88 | Nelarabine | Antimetabolite antineoplastic agent, used to treat T-cell acute lymphoblastic leukemia |  |
| 5366 | 1.81 | Noscapine | Antitussive agent |  |
| 279836 | 1.76 | Mitoxantrone | Topoisomerase inhibitor; antineoplastic agent | HIV<sup>43</sup>, VACV<sup>44</sup>,<br>HCV<sup>45</sup>, HSV<sup>46</sup>,<br>RVFV<sup>16</sup> |
| 11440 | 1.64 | Protopine | Calcium channel blocker and anti-inflammatory agent |  |
| 11926 | 1.60 | Tardolyt | Carcinogenic molecule from birthwort plants | HSV<sup>47</sup> |

HIV, Human immunodeficiency virus; HRV, human rhinovirus; HBV, hepatitis B virus; EBV, Epstein-Barr virus; EV, enterovirus; SARS-CoV, severe acute respiratory syndrome coronavirus; RSV, respiratory syncytial virus; PV, poliovirus; MV, measles virus; HSV, herpes simplex virus; HAV, hepatitis A virus; VACV, vaccinia virus

**Figure 1.**
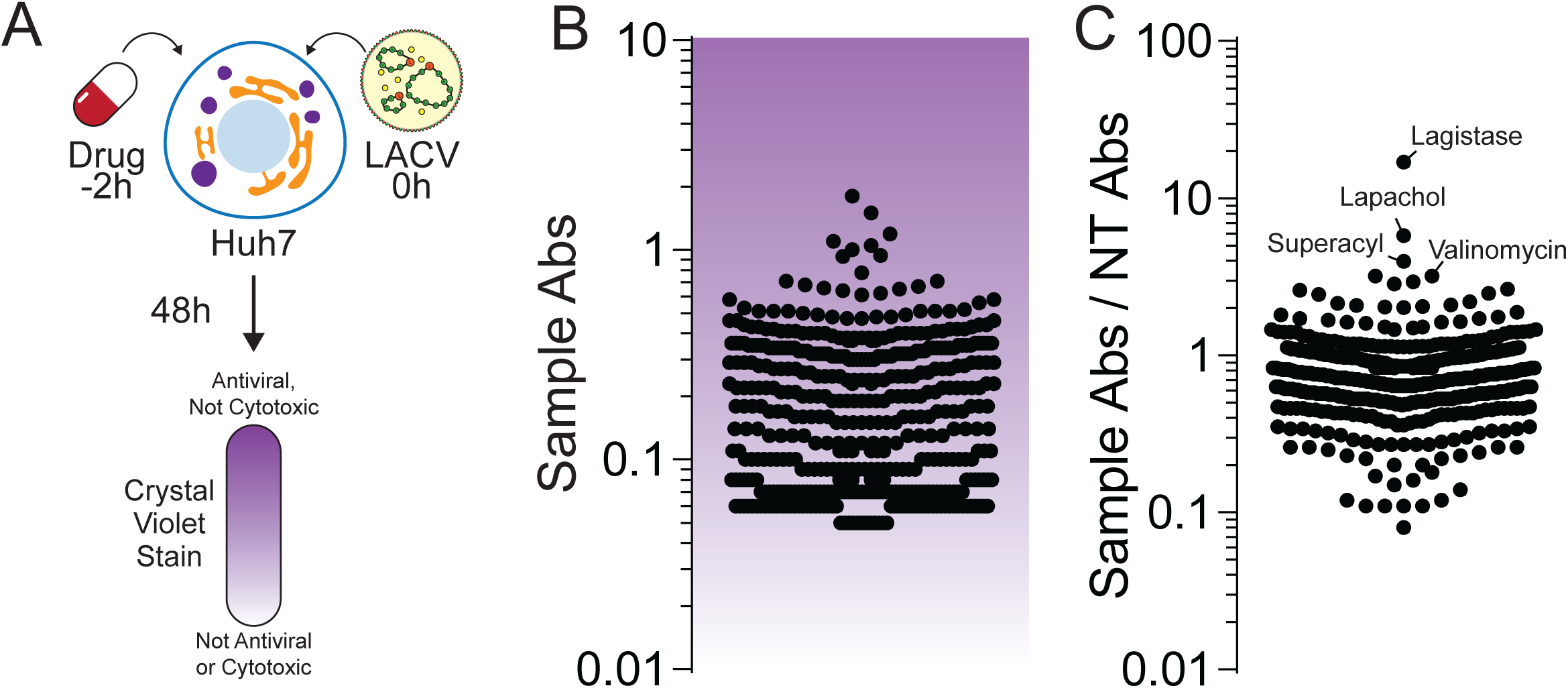
Rapid screening for LACV antivirals. Schematic of screen performed in these studies. (A) Potential antivirals were added to cells two hours prior to infection with La Crosse virus at MOI 0.01. At 48 hpi, cells were fixed and stained with crystal violet stain, which was then quantified, with darker staining cells surviving drug treatment and virus infection. (B) Quantification of antiviral activity. Each dot represents a single compound analyzed in our assay. (C) The quantification of antiviral activity as represented in (B) was compared to control cells that were not treated (Sample Abs / NT Abs) to obtain the relative antiviral activity. Top hits in this screen are called out.

### Valinomycin restricts LACV replication

We focused on valinomycin, as the molecule was a prominent hit in our screen, had a known mechanism relevant to LACV infection, and was not previously described to have antiviral activity against LACV. As an initial consideration of its antiviral activity, we performed secondary screening on Huh7 cells. Cells were seeded to confluency, treated with increasing doses of valinomycin, from 1 to 64 µM, and infected at MOI 0.1. At 48h, cells were fixed and stained with crystal violet, and stain was quantified by absorbance reading. We observed that valinomycin exhibited antiviral activity at doses above 10 µM, as crystal violet staining was stronger, suggesting more surviving cells (Figure 2A). Doses as high as 64 µM did not affect crystal violet stain, suggesting that cellular viability was not compromised. To confirm this phenotype with titers, we treated cells with increasing doses of valinomycin 2h prior to infection at MOI 0.1 and measured titers by plaque assay at 48 hpi. We observed that viral titers were significantly decreased compared to untreated controls (Figure 2B, dotted line) at concentrations above 1 µM. In fact, viral titers were reduced over 100-fold at 10 µM. We calculated an IC50 value of 1.4 µM. To confirm that cellular viability was not compromised, we used a fluorescent assay to measure cellular ATP content after treatment with increasing doses of valinomycin. We observed cellular toxicity at doses at and above 16 µM (Figure 2C, CC50 value of 14 µM), though no toxicity was observed either by cellular morphology or fluorescent ATP assay below 10 µM.

**Figure 2.**
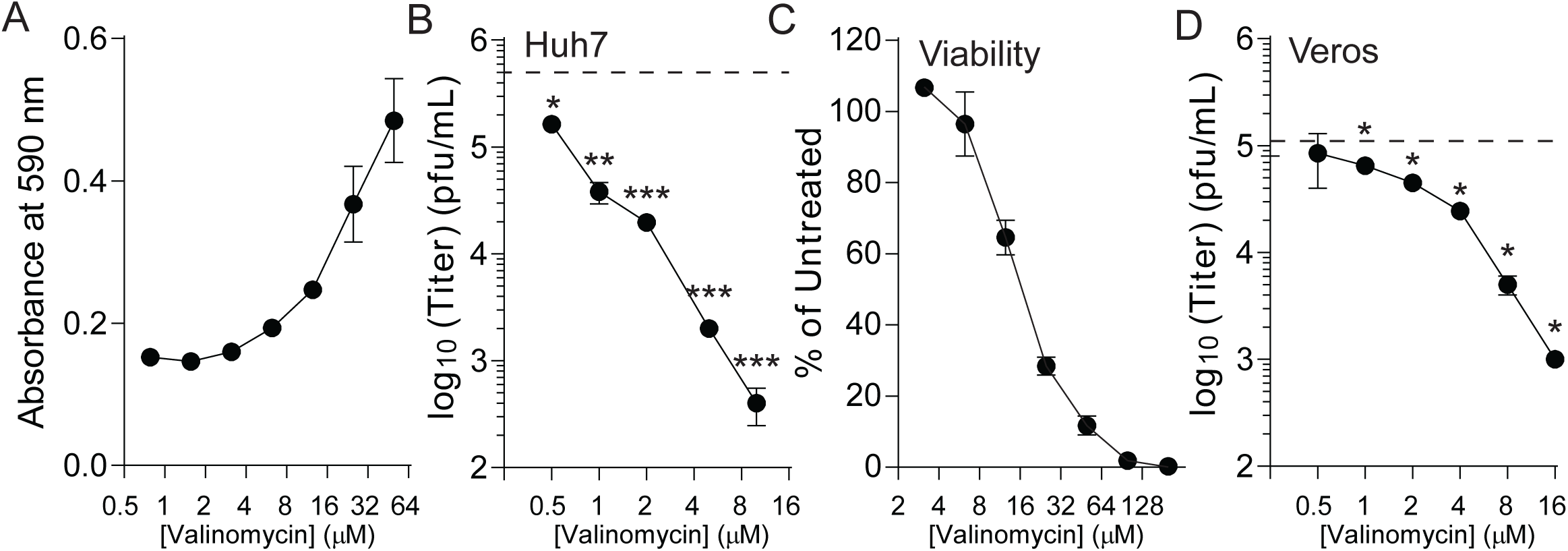
Valinomycin is antiviral. Huh7 cells were treated with increasing doses of valinomycin for two hours prior to LACV infection. (A) At 48 hpi, cells were stained with crystal violet and quantified. (B) Viral titers were measured by plaque assay. Dotted line indicates titer of untreated controls. (C) Viability was measured by fluorescent intracellular ATP assay. (D) Vero cells were treated and infected as Huh7 cells were. Viral titers were determined at 48 hpi. *p<0.05, **p<0.01, ***p<0.0001 using two-tailed Student’s T-test, n=3. Error bars represent one standard error of the mean.

As further confirmation of valinomycin’s antiviral activity, we measured cell-associated viral genomes. Huh7 cells were treated and infected as above and cell-associated RNA was collected at 48 hpi. RNA was purified, reverse transcribed, and analyzed via qPCR for Small, Medium, and Large genome segments, normalizing to cellular β-actin. Paralleling our titer data, valinomycin treatment reduced viral genome content in a dose-dependent manner (Figure 2D). Viral genome content was reduced upwards of 100-fold with 10 µM valinomycin treatment. To extend these results to other cell types, we treated and infected Vero-E6 cells as above and determined viral titers by plaque assay at 48 hpi. Again, we observed a significant reduction in viral titer with an IC50 value of 900 nM. In sum, our data suggest that valinomycin is antiviral at non-cytotoxic doses in multiple cell types, reducing both viral titers and cell-associated viral genomes in a dose-dependent manner.

### Valinomycin is antiviral over multiple rounds of infection

Our initial assays were performed at low MOI and viral titer measured at 48 hpi. To determine if valinomycin was antiviral over several rounds of replication, we treated Huh7 cells with 2 µM valinomycin two hours prior to infection at MOI 0.1 and subsequently collected samples to titer every 8h for 56h total. We found that LACV titers were significantly reduced at all times after 8 hpi (Figure 3A); in fact, virus failed to replicate above input virus titers (0h). To confirm that valinomycin was reducing virus replication, we measured viral RNA genomes. To this end, we treated cells with increasing doses of valinomycin, infected with LACV and collected cell-associated RNA in Trizol at 24 hpi. After purifying and reverse-transcribing, we performed qPCR using primers specific to the small, medium, and large genome segments. We observed that treatment with valinomycin significantly reduced the number of viral genomes by >90% with treatment and that no individual genome segment was affected more than another (Figure 3B). Together, these data suggest that valinomycin blocks virus replication and reduces viral RNA accumulation.

**Figure 3.**
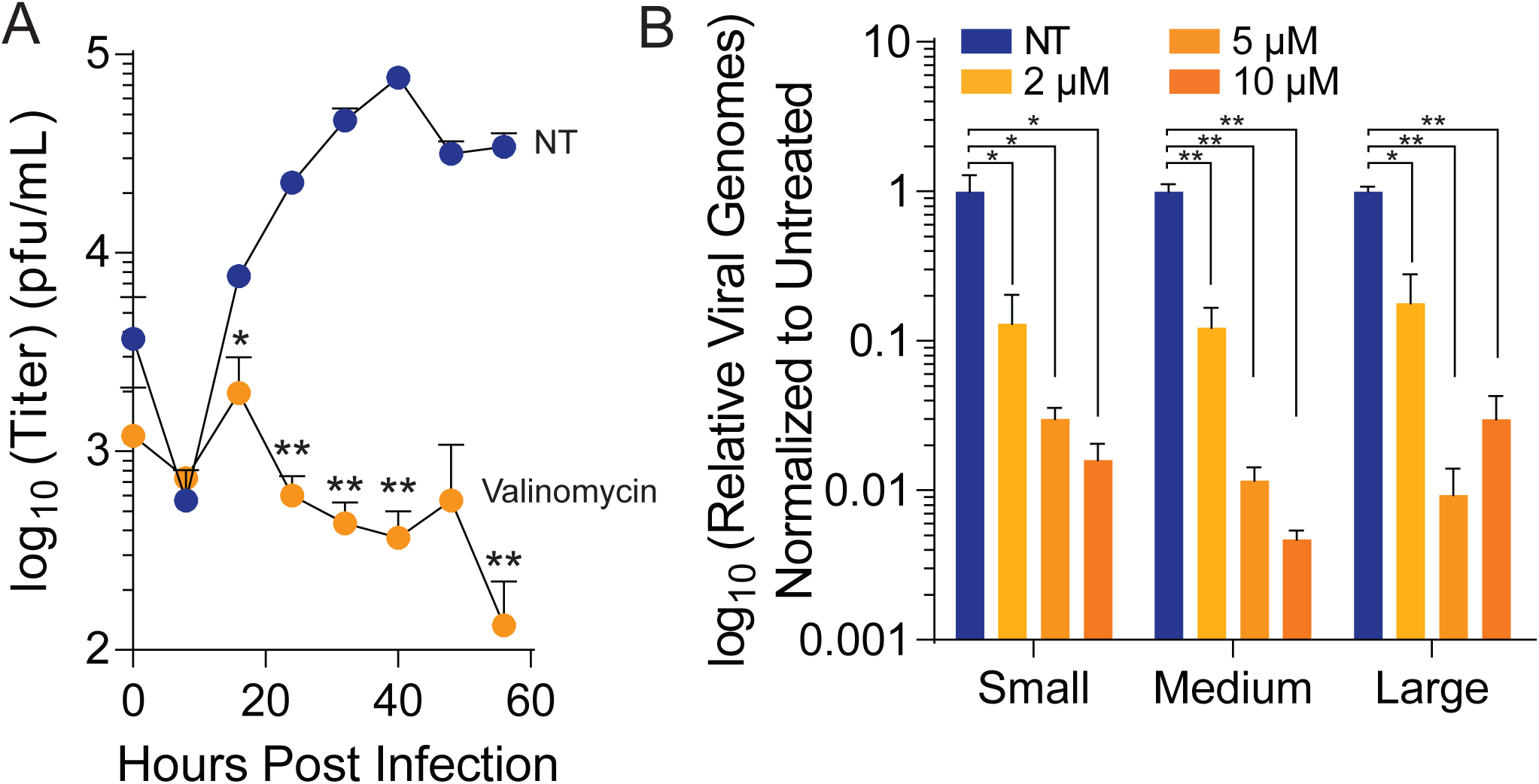
Valinomycin is antiviral over multiple rounds of infection. Huh7 cells were treated with 2 µM valinomycin for two hours prior to infection at MOI 0.01. (A) Cellular supernatant was collected at the times indicated and viral titers were determined by plaque assay. (B). Viral RNA was purified from infected cell supernatant at 48 hpi and quantified by qPCR using primers specific to each genome segment. *p<0.05, **p<0.01, ***p<0.001 using two-tailed Student’s T-test, n=2-3. Error bars represent one standard error of the mean.

### Valinomycin does not reduce viral particle infectivity

Because we observe significant reductions in LACV titers with valinomycin treatment, we hypothesized that valinomycin might be directly affecting cellular processes to reduce virus infection. Nonetheless, valinomycin is a cyclic peptide and could potentially directly inactivate viral particles, as seen previously^20^. To test whether valinomycin directly reduced virus infectivity, we directly incubated LACV with 2 µM valinomycin for 24h and directly titered the surviving virus at regular intervals. We found that valinomycin did not significantly alter viral titer over the time examined (Figure 4A), suggesting that valinomycin is not directly inactivating viral particles. We further examined the capacity of valinomycin to inactivate particles by incubating with increasing doses, up to 10 µM valinomycin, for 24h prior to directly titering. As in our timecourse, we observed no significant change in viral titers at any dose (Figure 4B), again suggesting that valinomycin does not directly inactivate LACV particles. As final confirmation of this phenotype, we measured viral RNA in viruses exposed to increasing doses of valinomycin. We then compared the relative number of genomes to the titer to calculate the genome-to-PFU ratio. We observed that this number did not change with valinomycin treatment, suggesting no change in specific infectivity (Figure 4C). In sum, these data suggest that valinomycin does not affect virus infectivity by directly acting on the virion.

**Figure 4.**
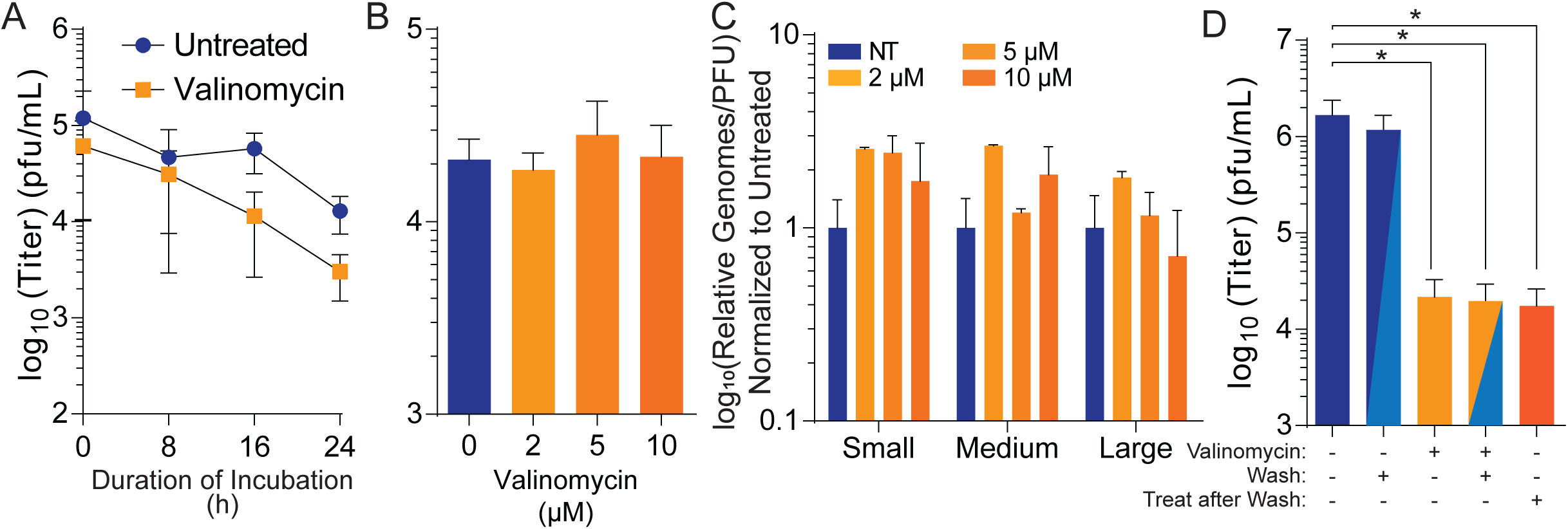
Valinomycin does not directly reduce particle infectivity. (A) Stock LACV was incubated with 2 µM valinomycin at 37°C for the times indicated before titering by plaque assay. (B) Stock virus was incubated with increasing does of valinomycin at 37°C for 24h prior to titering. (C) Viral genomes from (B) were quantified from purified RNA and compared to viral titers to calculate the relative viral genomes compared to infectious virus (PFU). (D) Huh7 cells were treated with 2 µM valinomycin and subsequently washed with PBS and replenished with fresh media as indicated. Fresh valinomycin was added as indicated following a PBS wash and media replenishment. *p<0.05, **p<0.01, ***p<0.001 using two-tailed Student’s T-test, n≥2. Error bars represent one standard error of the mean.

### Valinomycin activity alters host cell activity

We thus hypothesized that valinomycin’s antiviral activity was due to its effect on the cell. The role of potassium in bunyavirus infection has been well-documented, and bunyavirus entry is potassium dependent^21,22^. To test if valinomycin was affecting the cell rather than the virus, we treated Huh7 cells with 2 µM valinomycin and, immediately before infection, we washed away the drug. As a control, we maintained valinomycin on cells or replaced the valinomycin after washing away the initial treatment. We observed that even after removing and washing valinomycin from the cells, the antiviral activity persisted, as viral titers remained reduced to the same level as when valinomycin treatment is concurrent with infection (Figure 4D). Together, these data suggest that valinomycin does not directly inactivate viral particles but that treatment of cells reduces LACV infection, potentially by disrupting potassium-dependent entry.

### Ionophores are selectively antiviral

Given that valinomycin is a potassium ionophore, we wished to investigate whether other ionophores, for potassium or otherwise, were antiviral. Potassium ions play a crucial role in cellular entry; however, a role for sodium or calcium ions is not as well described. Marituba virus (MTBV) infection results in a sodium ion influx ^23^, but the origin or function of these ionic changes are not known. To determine if other ionophores might exhibit antiviral activity, we treated cells with increasing doses of nonactin (potassium and sodium ionophore), nigericin (hydrogen and potassium ionophore), calcium ionophore I, and sodium ionophore III (Figure 5A). Two hours later, we infected with LACV at MOI 0.01 and measured viral titers at 48 hpi. Both nonactin and nigericin exhibited significant antiviral activity, and viral titers were not measurable above 4 and 1 µM, respectively (Figure 5B). Treatment with sodium ionophore III resulted in a dose-dependent decrease in viral titers, and virus was not recovered above 10 µM. Interestingly, treatment with calcium ionophore I showed no changes in viral titer, even at the highest dose. Thus, we observe that LACV replication is disrupted by several ionophores, especially potassium ionophores, highlighting the role for potassium in virus infection.

**Figure 5.**
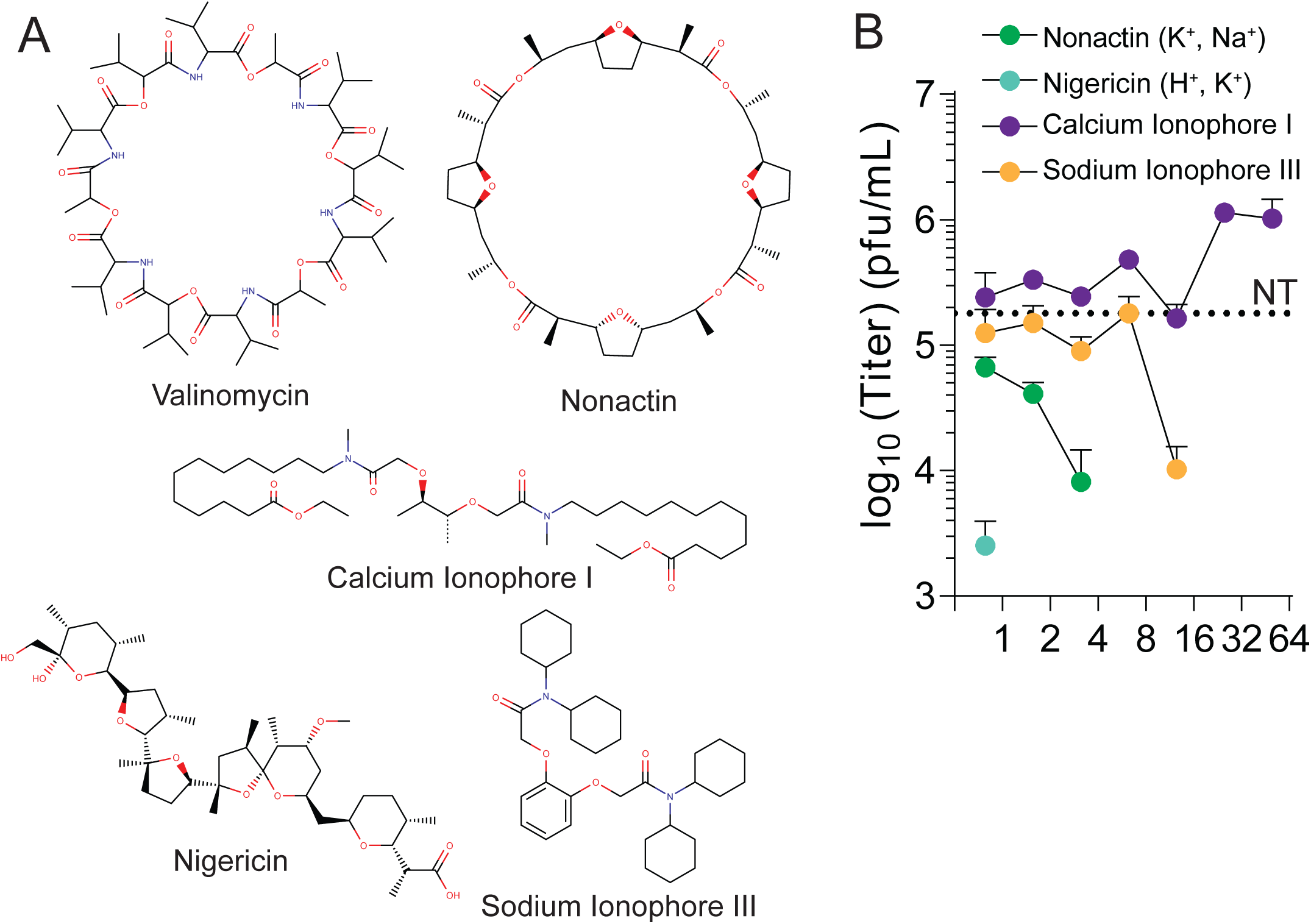
Ionophores are selectively antiviral. (A) Structures of valinomycin, nonactin, calcium ionophore I, nigericin, and sodium ionophore III. (B) Huh7 cells were treated with increasing doses of the iophores for two hours prior to LACV infection. Virus titers were determined at 48 hpi by plaque assay. Dotted line indicates titers from untreated (NT) control cells. N=3. Error bars represent one standard error of the mean.

### Valinomycin is broadly antiviral

LACV is related to several other medically-relevant bunyaviruses, including Keystone virus and Rift Valley fever virus. To determine whether these viruses respond to valinomycin treatment, we treated and infected Huh7 cells with these viruses and measured viral titers at 48 hpi. As with LACV, we observed a dose-dependent decrease in viral titers, with titers decreasing greater than 100-fold at concentrations above 2 µM (Figure 6A). To expand beyond bunyaviruses, we performed the same analysis with Zika virus (flavivirus) infection (Figure 6B), Coxsackievirus B3 and human rhinovirus 2 (picornaviruses) (Figure 6C), and SARS and MERS coronaviruses (coronaviruses) (Figure 6D). In all cases, valinomycin significantly reduced viral titers. Zika virus titers were most sensitive to valinomycin treatment, and virus was not recovered above 500 nM valinomycin treatment. Similarly, valinomycin treatment significantly disrupted both 229E-CoV and MERS-CoV infection. Thus, valinomycin exhibits broad antiviral activity, highlighting cellular potassium as a conserved and crucial host factor in virus replication.

**Figure 6.**
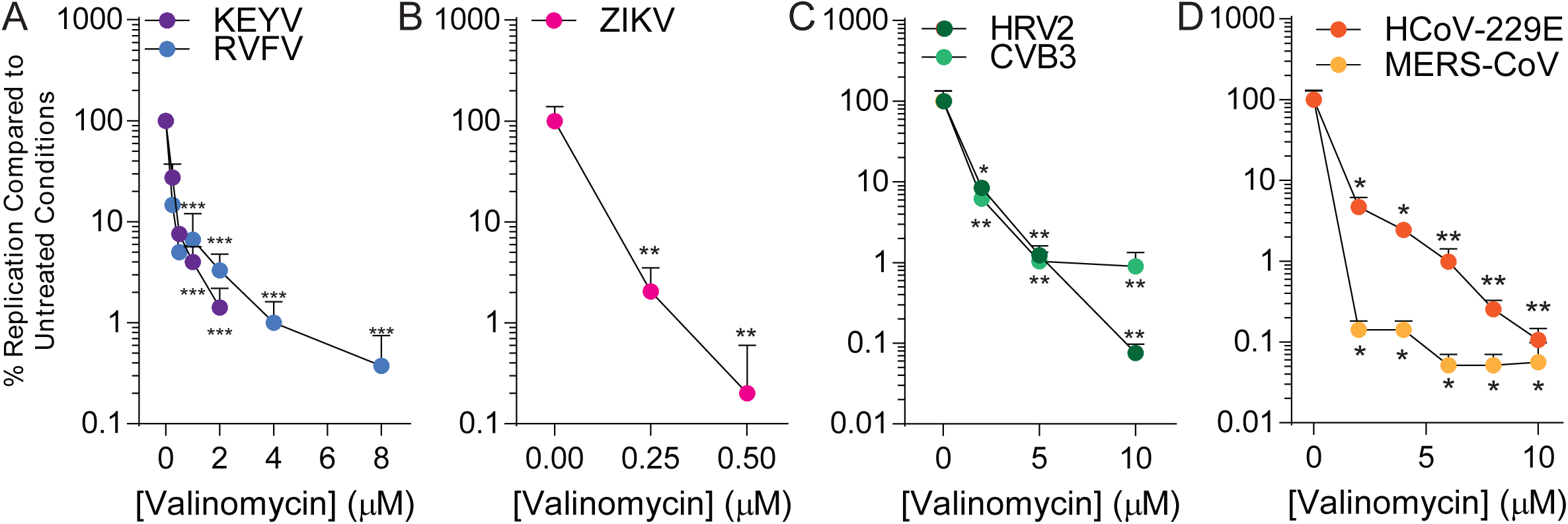
Valinomycin is broadly antiviral. Huh7 cells were treated with increasing doses of valinomycin for two hours prior to infection at MOI 0.1 with (A) bunyaviruses KEYV and RVFV (strain MP12) for 48 h, (B) flavivirus ZIKV for 48h, (C) enteroviruses HRV2 and CVB3 for 24h, and (D) coronaviruses HCoV-229E and MERS-CoV for 24h. *p<0.05, **p<0.01, ***p<0.001 using two-tailed Student’s T-test, n≥2. Error bars represent one standard error of the mean.

## Discussion

Treating virus infected patients, including encephalitic patients, is limited in the scope of available therapeutics. La Crosse virus lacks any approved treatment, and antiviral development could benefit not only LACV-infected patients but perhaps patients infected with related bunyaviruses. Unfortunately, antiviral development is a nonlinear process that has provided no promising targets for this specific virus to date. Our screen is the first to investigate a large number (>500) of compounds in their ability to reduce LACV infection *in vitro*, and our hits demonstrate that several known antivirals may exhibit anti-LACV activity. Additionally, the identification of novel antivirals, such as valinomycin provides novel avenues of antiviral development.

In our screens, valinomycin consistently exhibited antiviral activity and was selected for follow-up experiments. Based on our results, valinomycin is a host-targeted antiviral with broadspectrum activity. In fact, our tests against eight distinct viruses and four diverse families of virus suggest (1) valinomycin has activity against evolutionarily-distant viruses and (2) these viruses rely on this conserved host process for efficient replication. Precisely why each virus is sensitive to valinomycin is likely virus-specific, though mechanisms for bunyaviruses have been described^22,24^. As a potassium ionophore, valinomycin disrupts potassium ion gradients, resulting in aberrant cellular events, including endocytosis. This potassium-dependent endocytosis is required for efficient bunyavirus entry^24^. Additionally, a study using Hazara virus, a distantly related bunyavirus, demonstrated that the viral fusion spike conformation was potassium-sensitive^21^. Thus, valinomycin’s antiviral activity fits with the prescribed role of potassium ions during virus infection, both highlighting the importance of this pathway for virus infection and its potential as an antiviral target.

Valinomycin is a naturally occurring cycling peptide, synthesized by *Streptomyces* species as an antibiotic^25^. In our assays, we did not observe significant cellular toxicity, as measured either by gross cellular morphology or by measuring cellular ATP levels, until doses above 10 µM, while our IC50 of 1.4 µM suggests there is a window of therapeutic potential. However, given the drug’s toxicity *in vivo*, modification of the valinomycin structure may tune toxicity while maintaining antiviral activity. Additionally, using other, less toxic potassium ionophores may similarly function to inhibit LACV infection, as we observed with nonactin and nigericin. For outbreak viruses like SARS CoV or ZIKV, however, limited valinomycin usage might be beneficial. Interestingly, valinomycin was identified in an earlier screen for anti-SARS-CoV molecules^26^, underscoring the antiviral potential of valinomycin despite its negative characteristics. Importantly, the emergence of a new group 2B CoV, nCoV-2019 signals the ongoing threat and need to rapidly respond to novel emergent virues. Additional *in vivo* testing, combined with medicinal chemistry approaches to structural modification, would be necessary prior to clinical use.

In addition to valinomycin, our screen identified several tantalizing antiviral candidates. In fact, several known antivirals were identified (summarized in Table 1). Dasatinib has previously been described to inhibit dengue virus and HIV infection^27,28^. Quininone has activity against enterovirus proteases as well as reverse transcriptases^29,30^. As mentioned previously, mitoxantrone was identified in a screen for anti-RVFV molecules^16^. Additionally, we identified a number of cancer therapeutics, targeting processes such as angiogenesis and topoisomerase, that show significant promise. Nonetheless, further verification of antiviral activity and investigation of the mechanisms of action is necessary. Regardless, the development of novel antivirals is crucial to combat virus infection and to respond to the possibility of rapid virus dissemination and evolution. LACV is a significant threat to human health, and continued development of novel antivirals may prove fruitful if the virus were to spread, as arboviruses are wont to do.

## Materials and Methods

### Cell culture

Cells were maintained at 37°C in 5% CO_2_, in Dulbecco’s modified Eagle’s medium (DMEM; Life Technologies) with bovine serum and penicillin-streptomycin. Vero cells (BEI Resources) were supplemented with 10% new-born calf serum (NBCS; Thermo-Fischer) and Huh7 cells, kindly provided by Dr. Susan Uprichard, were supplemented with 10% fetal bovine serum (FBS; Thermo-Fischer).

### Drug treatment

For standard treatment experiments Huh7 cells were infected at a multiplicity of infection (MOI) of 0.01 PFU/cell, unless otherwise indicated, with LACV, KEYV, RVFV MP-12, ZIKV and concurrently treated with ionophores (valinomycin, nigericin, nonactin, calcium ionophore I, sodium ionophore III; Cayman Chemical) dissolved in DMSO. Cells were then incubated at 37°C in 5% CO_2_ for 48 hours. For direct incubation experiments 100 µL of LACV stock virus was incubated with increasing concentrations of valinomycin over various time periods at 37°C. For wash away experiments cells were seeded as stated above for standard treatment experiments. Cells were treated four hours prior to removing media and washing with phosphate buffer saline (PBS). Media containing LACV was then replaced and cells were incubated for 48 hours. Chemical structures were recreated using MarvinSketch 19.26 (ChemAxon Ltd.).

### Rapid screening antiviral compounds

Huh7 were seeded in 96 well plates, infected with LACV at a MOI of 0.01 and concurrently treated with 100 µM of each compound from the NIH DTP compound plates. Cells were incubated at 37°C in 5% CO_2_ for 48 hours. Media was aspirated and cells were fixed with 4% formalin and live cells were stained with crystal violet solution (Sigma-Aldrich). Excess stain was removed in a mild bleach solution and allowed to dry for 24h. Remaining crystal violet stain was resuspended in 10% acetic acid. Absorbance at 590 nm was detected using a BioTek Synergy H1 plate reader.

### Infection and enumeration of viral titers

RVFV MP-12^31^, LACV, and KEYV were derived from the first passage of virus in Huh7 cells. ZIKV (MR766) was derived from the first passage of virus in Vero cells. CVB3 (Nancy strain) and HRV2 were derived from the first passage of virus in HeLa cells. ZIKV, LACV, and, KEYV were obtained from Biodefense and Emerging Infections (BEI) Research Resources. HCoV-229E and MERS-CoV were propagated and quantitated via standard methods as previously described^20^.For all infections, drug was maintained throughout infection as designated. Viral stocks were maintained at −80°C. For infection, virus was diluted in serum-free DMEM for a multiplicity of infection (MOI) of 0.1 on Huh7 cells, unless otherwise indicated. Viral inoculum was overlain on cells for 10 to 30 minutes, and the cells were washed with PBS before replenishment of media. Supernatants were collected at the times specified. Dilutions of cell supernatant were prepared in serum-free DMEM and used to inoculate confluent monolayer of Vero cells for 10 to 15 min at 37°C. Cells were overlain with 0.8% agarose in DMEM containing 2% NBCS. CVB3 and HRV2 samples incubated for 2 days, RVFV MP-12, ZIKV, and LACV samples incubated for 4 days and KEYV samples incubated for 5 days at 37°C. Following appropriate incubation, cells were fixed with 4% formalin and revealed with crystal violet solution (10% crystal violet; Sigma-Aldrich). Plaques were enumerated and used to back-calculate the number of plaque forming units (pfu) per milliliter of collected volume.

### RNA purification and cDNA synthesis

Media was cleared from cells and Trizol reagent (Zymo Research) directly added. Lysate was then collected, and RNA was purified according to the manufacturer’s protocol utilizing the Direct-zol RNA Miniprep Plus Kit (Zymo Research). Purified RNA was subsequently used for cDNA synthesis using High Capacity cDNA Reverse Transcription Kits (Thermo-Fischer), according to the manufacturer’s protocol, with 10-100 ng of RNA and random hexamer primers.

### Viral genome quantification

Following cDNA synthesis, qRT-PCR was performed using the QuantStudio3 (Applied Biosystems by Thermo-Fischer) and SYBR green mastermix (DotScientific). Samples were held at 95°C for 2 mins prior to 40 cycles of 95°C for 1s and 60°C for 30s. Primers were verified for linearity using eight-fold serial diluted cDNA and checked for specificity via melt curve analysis following by agarose gel electrophoresis. All samples were used to normalize to total RNA using the ΔC_T_ method.

### Genome-to-PFU ratio calculations

The number of viral genomes quantified as described above were divided by the viral titer, as determined by plaque assay, to measure the genome-to-PFU ratio. Values obtained were normalized to untreated conditions to obtain the relative genome-to-PFU ratio. Primers used were: Small 5’-GGC-AGG-TGG-AGG-TTA-TCA-AT-3’ (forward), 5’-AAG-GAC-CCA-TCT-GGC-TAA-ATA-C-3’ (reverse, Medium 5’-CCT-GCC-TAG-AGA-CTG-AGA-GTA-T-3’ (forward), 5’-GAG-TTG-CAA-TGT-TGG-TGT-AAG-G-3’ (reverse), Large 5’-ACT-GGA-AGG-TCG-AGG-ATC-TAA-3’ (forward), 5’-GTC-GCT-TGT-CTC-ACC-CAT-AAT-A-3’ (reverse), GAPDH 5’-GAT-TCC-ACC-CAT-GGC-AAA-TTC-3’ (forward), 5’-CTG-GAA-GAT-GGT-GAT-GGG-ATT-3’ (reverse).

### Statistical Analysis

Prism 6 (GraphPad) was used to generate graphs and perform statistical analysis. For all analyses, one-tailed Student’s t test was used to compare groups, unless otherwise noted, with a = 0.05. For tests of sample proportions, p values were derived from calculated Z scores with two tails and a = 0.05.

## Acknowledgments

We are gracious to Drs. Susan Uprichard, Justin Harbison, and Gail Reid for critical discussion of the data and manuscript. We also thank Susan Uprichard for Huh7 cells, Shinji Makino and Kaori Terasaki for generously providing the MP-12 strain of RVFV, and Bill Jackson for HRV2. Research was supported by grant from NIAID of the NIH (U19AI100625 to VDM). Research was also supported by STARs Award provided by the University of Texas System to VDM and trainee funding provided by the Graduate School of Biomedical Sciences at UTMB.

